# Distinct roles of SNR, speech Intelligibility, and attentional effort on neural speech tracking in noise

**DOI:** 10.1101/2024.10.10.616515

**Authors:** Xiaomin He, Vinay S Raghavan, Nima Mesgarani

## Abstract

Robust neural encoding of speech in noise is influenced by several factors, including signal-to-noise ratio (SNR), speech intelligibility (SI), and attentional effort (AE). Yet, the interaction and distinct role of these factors remain unclear. In this study, fourteen native English speakers performed selective speech listening tasks at various SNR levels while EEG responses were recorded. Attentional performance was assessed using a repeated word detection task, and attentional effort was inferred from subjects’ gaze velocity. Results indicate that both SNR and SI enhance neural tracking of target speech, with distinct effects influenced by the previously overlooked role of attentional effort. Specifically, at high levels of SI, increasing SNR leads to reduced attentional effort, which in turn decreases neural speech tracking. Our findings highlight the importance of differentiating the roles of SNR, SI, and AE in neural speech processing and advance our understanding of how noisy speech is processed in the auditory pathway.

## 1. Introduction

The neural encoding of speech in noise is an essential process that enables speech comprehension in complex auditory scenes. Various objective and subjective factors influence how the auditory cortex processes noisy speech. Objective factors include the signal-to-noise ratio (SNR), representing the physical properties of the acoustic signal and its masking by background noise. Speech intelligibility (SI), on the other hand, is a subjective measure that reflects the listener’s ability to recognize spoken words and depends not only on SNR but also on the listener’s auditory processing capabilities (Nilsson et al., 1994; Sharma et al., 2013). Attentional performance (AP) is another subjective factor that pertains to the listener’s ability to selectively concentrate on one speech stream among many and filter out unwanted sounds in complex auditory scenes. Another related yet distinct factor is attentional effort (AE) (Sarter et al., 2006), which involves the cognitive resources expended to focus on the attended talker while ignoring distractions and is influenced by listener’s engagement, fatigue, and overall task difficulty (Bruya and Tang, 2018; Sarter et al., 2006; Strauss and Francis, 2017). While these factors are interconnected, they are mechanistically distinct. SNR is an external, quantifiable measure, whereas intelligibility and attention are subjective experiences that vary across individuals, even in identical acoustic settings. This differentiation underscores the complexity of auditory processing and the gaps in our understanding of how these elements collectively influence neural speech encoding.

Past research has extensively studied the neural encoding of speech in noise, emphasizing the role of SNR and speech intelligibility. Studies demonstrated that increasing SNR generally enhances intelligibility and neural speech encoding (Das et al., 2018; Decruy et al., 2020a; Ding and Simon, 2013; Lesenfants et al., 2019; Vanthornhout et al., 2018). Others have used varying degrees of visual congruency to modulate intelligibility and examined its impact on neural speech encoding (Crosse et al., 2015; Iotzov and Parra, 2019). Further work has identified different response components that differentially reflect SNR or intelligibility, such as frequency bands (Etard and Reichenbach, 2019; Vanthornhout et al., 2018), temporal components (Decruy et al., 2020a; Ding and Simon, 2013; Yasmin et al., 2023), and response latency (Yasmin et al., 2023). However, these findings often imply a monotonic relationship between SNR, intelligibility, and neural encoding, which oversimplifies the dynamic interaction among these features (Krueger et al., 2017). For instance, increasing noise levels under specific conditions can enhance neural tracking (Das et al., 2018; Lesenfants et al., 2019), and higher intelligibility does not always correlate with increased neural encoding (Etard and Reichenbach, 2019). Moreover, past research typically used standard speech-in-noise tasks to measure intelligibility, often separating this assessment from the task used to evaluate neural responses. Such intelligibility tasks typically involve asking subjects to repeat short sentences heard in noisy environments (Feng and Chen, 2022; Nilsson et al., 1994; Sharma et al., 2013), a method that may only partially capture the complexities of real-world listening due to its limited engagement with challenging factors such as attention span and effort. It has been shown that attentional performance significantly modulates neural speech encoding (Ding and Simon, 2013; Mesgarani and Chang, 2012; O’Sullivan et al., 2015), where SNR can considerably change target speech intelligibility (Brungart et al., 2001) and the attentional effort required to maintain focus on a talker (Cui and Herrmann, 2023; Dimitrijevic et al., 2019; Zekveld et al., 2006). These findings suggest a complex interplay between SNR, intelligibility, attention, and their impact on neural speech encoding (Devocht et al., 2017; Krueger et al., 2017; Yasmin et al., 2023), highlighting a critical gap in our holistic understanding of how these factors individually and collectively shape neural encoding of speech in noise.

Our study aims to address the need for a comprehensive analysis integrating these dimensions (SNR, SI, and AE) to fully elucidate their combined impact on neural encoding. We examined neural responses to speech in noise through a multifaceted approach incorporating a high-resolution range of SNR values. We used a repeated word detection task (Kirchner, 1958; Laffere et al., 2020; Marinato and Baldauf, 2019), designed to continually monitor subjects’ behavior in a manner that integrates attentional performance with the assessment of speech intelligibility, allowing us to capture the variability of attentional engagement in natural listening conditions. Additionally, we estimated attentional effort by analyzing gaze velocity (Ala et al., 2020; Ciccarelli et al., 2019; Gopher, 1973) to understand their collective impact on EEG signals. Our findings advance our holistic understanding of noisy speech processing in the auditory cortex and have practical implications for designing auditory technologies to improve speech perception under challenging listening conditions.

## 2. Materials and Methods

### 2.1. Participants

Fourteen native American English speakers (7 males; mean ± standard deviation (SD) age, 24.86 ± 4.4 years) with self-reported normal hearing participated in the experiment. The study followed the protocol approved by the Institutional Review Board of Columbia University (Protocol Number: AAAR7230). Participants were paid for their time as well as a bonus based on their task performance (1-back detection hit rate).

### 2.2. Experiment Procedures

#### 2.2.1. Experiment 1: Measuring Intelligibility by Connected Speech Test

Speech intelligibility (SI) was measured with the Connected Speech Test (CST) (Cox et al., 1987) in experiment 1. Subjects listened to a series of connected short sentences from daily familiar topics with one sentence at a time. The sentences were normalized to 65dB and were covered with noises at different SNRs ranging from -12 dB to 4 dB. Subjects were asked to verbally repeat the words they heard. Stimuli were synthesized by Google Text-To-Speech API (WaveNet) (Oord et al., 2016) with four different voices (2 males and 2 females), and played by two loudspeakers placed at ± 45 degrees. Experimenters recorded subjects’ responding accuracy and regressed for individual SI curves afterward.

#### 2.2.2 Experiment 2: Multi-talker Speech-in-Noise Perception Test

American English podcast stories were synthesized by Google Text-to-Speech API with the same setting as experiment 1. 160 trials of context-continuous stories (average length ∼35s) were also played by two loudspeakers placed to ± 45 degrees of subjects (Fig 1A). During each trial, subjects were presented with two speech streams covered by naturalistic background noises (babble or street noise). The target speech was normalized to 65 dB, and its SNR ranged from - 12 dB to 4 dB.

**Fig 1.**
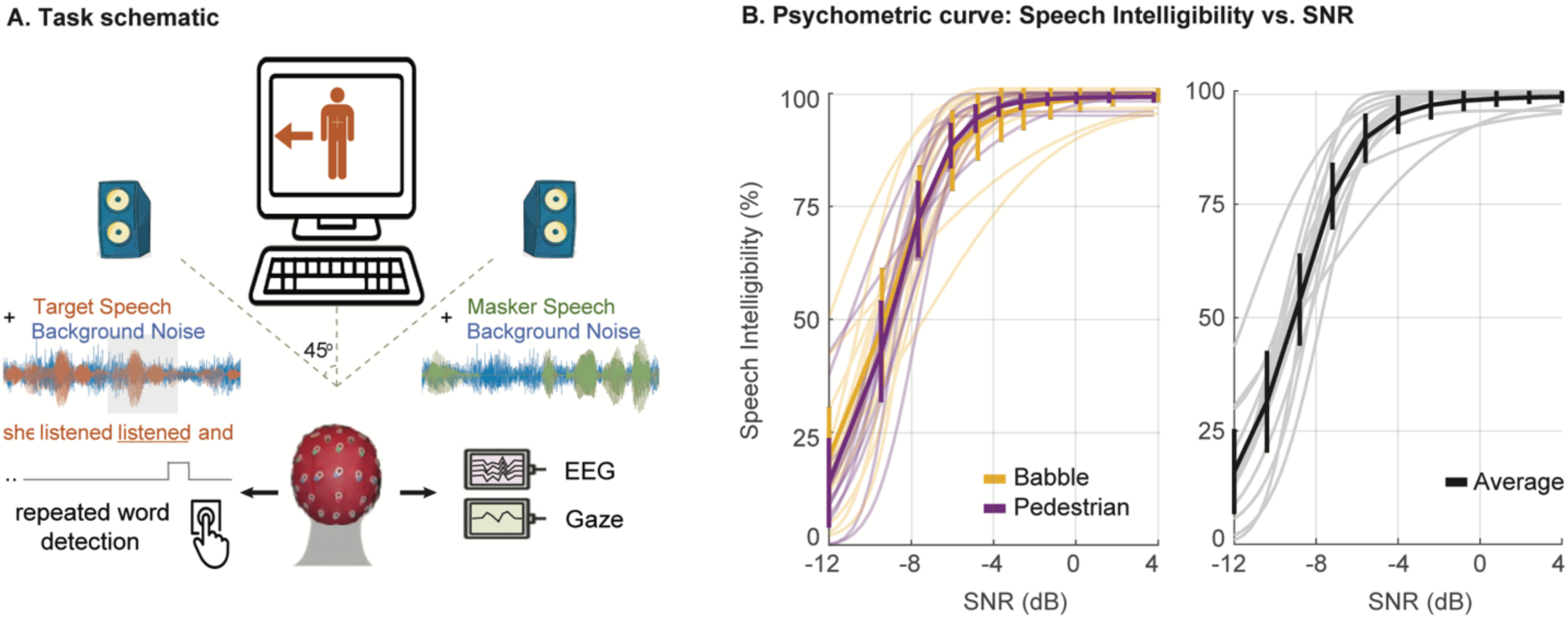
Illustration of the task schematic and psychometric curves. (**A**) The general task schematic. Subjects were instructed to focus on the target talker while ignoring the masker talker and the background noise. EEG, gaze activities, and button press were recorded while subjects performed the tasks. The subjects press a buzzer when they hear a repeated word in the target stream. (**B**) Speech intelligibility (SI) is measured using a connected speech task to derive psychometric curves for pedestrian and babble noises. No significant difference appears between the noise types.

Subjects were instructed to focus on the speaker whose gender and direction were specified by the icon on the monitor. Three repeated words were inserted in both speech streams. For simplicity, we selected semantically important keywords as repeated words, excluding articles, prepositions, and conjunctions. During the experiment, subjects needed to press the buzzer whenever they captured a repeated word from the target speaker. After each round of 16 trials, experimenters calculated the buzzer responses to the repeated words (1-back detection hit rate) and reported to subjects as feedback. Experimenters also asked subjects to summarize the stories they heard briefly. However, only 1-back detection hit rate was evaluated as a basis for compensation, to avoid unnecessary memory load for subjects.

Subjects were also instructed to keep their gaze on the monitor and minimize head movement during each trial. To facilitate tracking of target speech in extremely adverse trials with low SNRs, a 3-second window was used at the start of each trial where masker speech and noise gradually increased to the pre-set SNR. These windows were removed in later analyses.

### 2.3. Data Acquisition and Preprocessing

In Experiment 1, the word recalling accuracy for each SNR bin was manually recorded for later regressing the *SNR-SI* psychometric curve (Fig 1B; Details also in **2.4.2 Speech Perceptual** Attributes).

In Experiment 2, buzzer responses to the repeated words, 64-channel EEG, and eye-tracking data were recorded for each trial. Among them, buzzer responses and EEG were recorded by g.HIAMP (g.tec, Australia). Eye tracking data were calibrated and acquired from Tobii Pro Nano (Tobii, Sweden). All data was streamed from Simulink (Mathworks, MA, USA) at 1200 Hz with a 60Hz notch. Afterward, EEG data were downsampled to 100Hz with an anti-aliasing filter. Channels with unusual standard deviations were automatically detected and replaced using spherical interpolation of the remaining channels (Delorme and Makeig, 2004; Kang et al., 2015; Perrin et al., 1989).

Speech envelopes for both target and masker speakers were firstly extracted by a nonlinear, iterative (NLI) method (Horwitz-Martin et al., 2016) and secondly downsampled to 100Hz to match with the EEG recordings. Finally, each envelope was z-scored to zero mean and unit variance.

Blink detection and gaze tracking were completed and preprocessed automatically by Tobii Pro SDK (Tobii, Sweden) with a sampling frequency of 60Hz. The gaze coordinates were normalized to (0,0) and (1,1) within the screen.

### 2.4. Measurement of objective and subjective features

#### 2.4.1. Speech objective attribute: Signal-to-Noise ratio (SNR) of target speech

In both experiments, the volume of target speeches was normalized to 65 dB. The masker speech and bi-channel noises were at equalized volume to form the SNRs distribution from -12 dB to 4 dB. The SNRs were computed in the following formula:

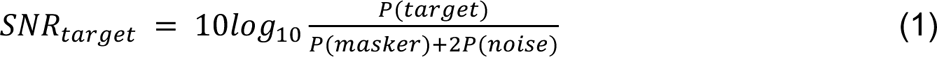

**P**: power of stimuli

The bi-channel noises used in this study for SNR adjustments are:

- The babble noise was 10-speaker babble derived from the AzBio test (Spahr et al., 2012).
- The street noise was the pedestrian area recording from CHiME3 (Barker et al., 2015) but with any salient interference removed (e.g. car horn, high-pitch car brake sound, intelligible pedestrians’ talking, etc.).

Noise audios were truncated as long as the formal trial (∼35s per trial).

#### 2.4.2. Speech perceptual attribute: Speech Intelligibility (SI)

Speech Intelligibility (SI) was measured by the Connected Speech Test (Cox et al., 1987). In Experiment 1, experimenters manually filed the subjects’ word recalling accuracy for each SNR bin in the range of -12 dB to 4 dB. Then, the psychometric curve between SNRs and word recall accuracy was fitted by *psignifit* toolbox, which implements the maximum-likelihood method described by (Wichmann and Hill, 2001a, 2001b) , and a customized logistic function:

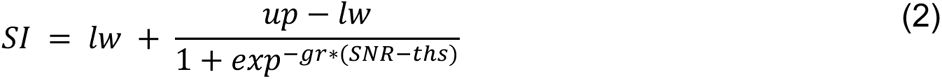

**lw:** lower bound (defined to approach 0); **up:** upper bound (defined to approach 1); **gr:** growth rate; **SNR:** signal-to-noise ratio, range from -12 to 4 dB; **ths:** threshold when SI = 50%, defined in the range of SNR.

Among the two approaches, the one producing a regressed curve with higher *R*^/^ and lower *RMSE* was selected. From the selected curve, the corresponding SI for each SNR in Experiment 2 was read.

#### 2.4.3. Attention Measures

- Attentional performance (AP): Single-trial 1-back detection hit rate (HR) As mentioned above, single-trial 1-back detection hit rate (HR), as a measure of subjects’ performance in terms of attention focus for each trial, was computed by the buzzer-hitting performance (Kirchner, 1958; Laffere et al., 2020; Marinato and Baldauf, 2019). There were 3 words inserted for each trial. Therefore, the range of HR was [0, ⅓, ⅔, 1]. Intuitively, high HR in a trial indicates a better attentional performance.
- Attentional effort (AE): Gaze Velocity (GV) Concentration periods are associated with suppressed irrelevant physiological activities, evidenced by reduced ocular movements (saccade and micro-saccade rate) and blink rates, as well as prolonged fixation (Abeles et al., 2020; Braga et al., 2016; Contadini-Wright et al., 2023; Cui and Herrmann, 2023). Oculomotor activity, with its anatomical overlap with the attention-related network (Corbetta et al., 1998), is, therefore, a valuable metric for evaluating attentional effort with superiority in stability across age (Bruenech, 2008), and the relationship has been justified by (Ala et al., 2020; Ciccarelli et al., 2019; Gopher, 1973). In this paper, we use averaged gaze velocity (GV) to quantify attentional effort for each trial, as it reflects the overall intensity of oculomotor activity, including saccade and micro-saccade. A higher GV, indicating more frequent oculomotor activity, suggests reduced attentional effort (Ala et al., 2020; Ciccarelli et al., 2019; Gopher, 1973). The gaze coordinates were recorded and normalized between (0,0) and (1,1) by the Tobii Pro Nano screen-based eye tracker. To calculate actual gaze angular velocity, we first restored relative coordinates to screen size, then computed and averaged the absolute value of the derivative of gaze coordinates over time within each trial. Finally, using the (approximately) 0.6m distance from the subject’s seat to the screen, we calculated gaze velocity (GV) using trigonometric functions (Diaz et al., 2013).

### 2.4. Attention Decoding

The classical approach for auditory attention decoding (AAD) is to model the linear projection between neural electrophysiological recordings and speech features (O’Sullivan et al., 2015), such as speech envelope. Once the model is trained, AAD correlates—the correlations between the speech features and their reconstruction from neural recordings—are compared for target and masker speech to decode auditory attention. Over and above the conventional forward or backward regularized linear model that applies transformation solely on one side of this projection (either neural signal or speech features), the Canonical Correlation Analysis (CCA) approach transforms both neural recordings and speech features for significantly better correlations scores (Dähne et al., 2015; de Cheveigné et al., 2018).

We adopt the CCA algorithms for target speech decoding. Speech envelope, which is the slow modulation of speech and proved to be feasible for neural speech tracking with EEG (Horton et al., 2014; Mesgarani et al., 2009; O’Sullivan et al., 2015), was extracted from both target and masker speech. Envelopes for clean speech and the multi-talker EEG recordings were first downsampled to 100Hz for a sampling rate match. Second, stimuli and neural recordings were windowed for overlapping receptive fields. Time-lagged matrices were produced for envelopes and EEG recordings. For EEG, the receptive fields were 400 ms and for speech envelope, the receptive fields were 200ms. Third, for each subject, subject-wise CCA-based linear models for both speeches were trained in a leave-one-out cross-validation setting.

The stimuli-response mapping is quantified by the trained model and evaluated by Pearson’s correlation between transformed stimuli and neural responses. As the target and masker stimuli are not identically encoded in the brain (Ding and Simon, 2012a), we computed this correlation for both the target and masker speech. The correlation for the target is referred to as rT, and for the masker speech is rM. We also defined rD as their difference (rD = rT-rM) to quantitively represent the different intensities of neural entrainment caused by selective attention. rD > 0 indicates a successful attention-decoded trial.

Moreover, to investigate the neural modulation pattern under varying speech conditions, we estimated temporal response functions (Ding and Simon, 2012b; Lalor et al., 2009) of target speech using regularized linear regression. This approach minimizes the mean-square error between the actual neural recordings and the predicted values. The training and prediction processes for each subject were also conducted in a leave-one-out cross-validation fashion using the mTRF toolbox (Crosse et al., 2016).

## 3. Result

Fourteen participants were instructed to perform selective listening tasks, focusing on a target speaker’s speech (attended stream) while ignoring a masker speaker (non-target, unattended) and background noises. We recorded 64-channel EEG, gaze velocity, and buzzer press responses to capture the participants’ neural and behavioral responses in real-time (Fig 1A). For each participant, speech intelligibility (SI) was measured using a connected speech test (Cox et al., 1987) prior to the actual experiment. As the difference in psychometric curves between different types of noise was negligible (Fig 1B, left), we adopted an average psychometric curve for each subject to streamline the analysis (Fig 1B, right).

### 3.1. Distinct Impacts of SNR and SI on Neural Speech Tracking: SI Enhances Tracking, While High SNR Reduces It

To measure the strength of neural tracking for target and masker speech, we trained CCA-based linear models to quantify neural speech tracking for each talker. We computed Pearson’s correlation between the transformed speech envelope and neural recordings to measure the strength of neural speech tracking for the target (rT) and masker (rM) speech. The difference in correlation between target and masker speech (rD *=* rT – rM) was used as a single measure to reflect how well participants followed the target speech while suppressing the masker speech.

To accurately assess the impact of SNR and SI on target and masker neural speech tracking, it is crucial to distinguish between these two highly correlated factors. We addressed this by analyzing their effect on neural speech tracking as a function of both SI and SNR. Our analysis revealed distinct patterns in how SNR and SI affect neural speech tracking. Fig 2A-2C show the averaged neural tracking correlations across subjects for target (rT) and masker speech (rM) and their difference (rD) for different SI and SNR values. We found that while increasing SNR and SI generally increase rT (enhanced neural tracking of the target speech) and decrease rM (suppressed neural tracking of the masker speech), this relationship shifts when SI is sufficiently high. Specifically, under high SI conditions (i.e., SI>80%), increasing SNR reduces rT and increases rM, indicating decreased neural tracking of the target speech while increasing the tracking of the masker speaker. The average plots across SI and SNR in Fig 2D and 2E further illustrate these effects, showing that SI has a nearly monotonic relationship with rT, rM, and rD, while SNR’s impact on these values reverses beyond approximately -1.6 dB. This indicates that in easier listening conditions, when the target talker is highly intelligible, increasing the SNRs of the masker talker can paradoxically reduce its neural speech tracking. In Fig 2F, we quantized SI into 6 bins from 0 to full intelligibility, each denoted by a different color. Trials categorized as ’Full SI’ are marked in black and represent instances where SI reached its plateau on individual psychometric curves. Fig 2G illustrates the linear relationship between SNR and rD within each SI bin, revealing a progression from positive to neutral to negative correlation between rD and SNR as SI increases. In summary, these results demonstrate that while SNR and SI are strongly correlated, they have distinct and sometimes opposing effects on neural speech tracking, which we will explore in greater detail in the next section

**Fig 2.**
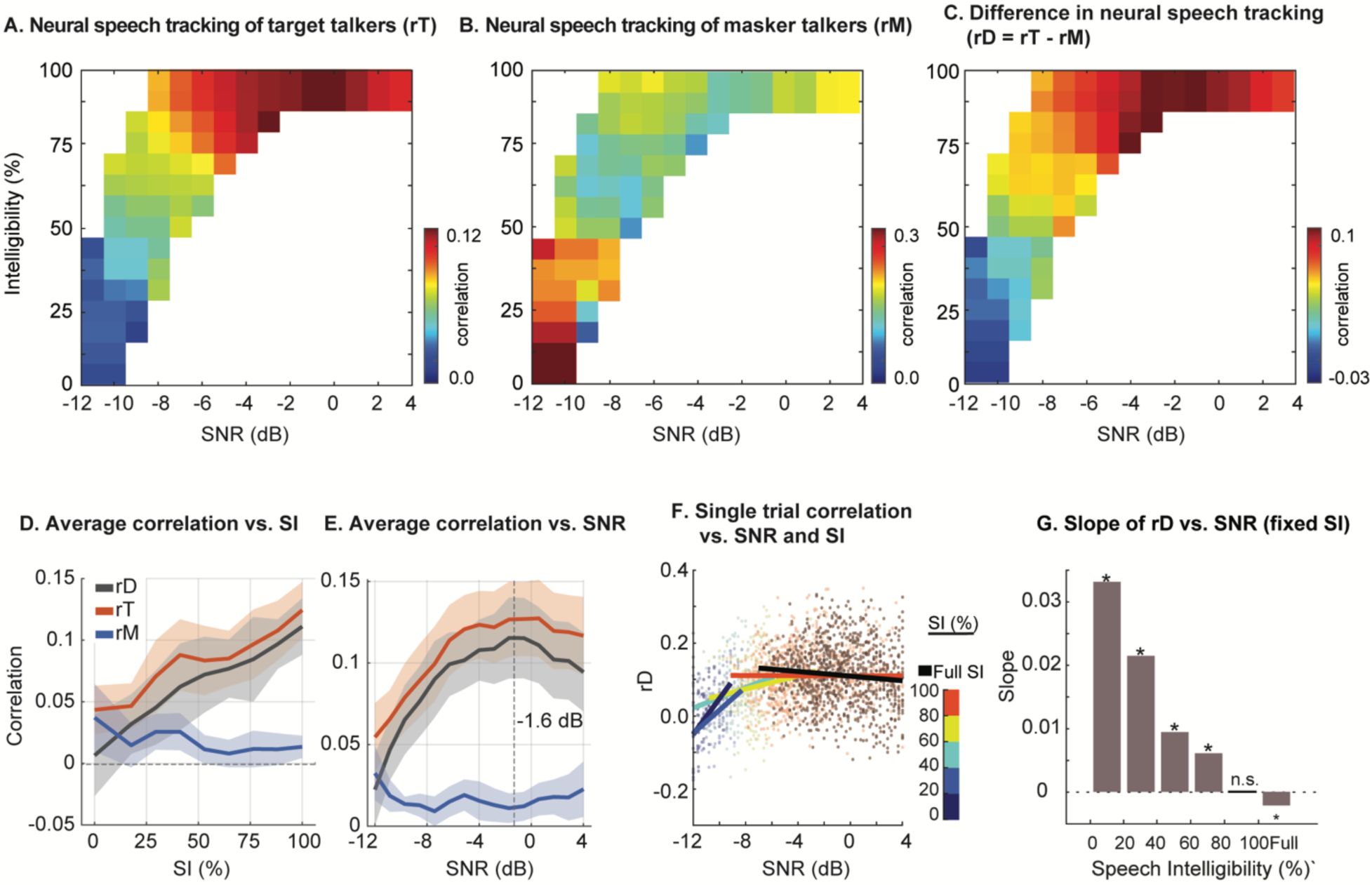
The relationship between neural speech tracking for target and masker speech with variables SNRs and SI. (**A**) rT: the correlation of the target neural speech tracking across different SNR and SI values; (**B**) rM: the correlation of masker neural speech tracking across different SNR and SI values; (**C**) rD: the difference between neural speech tracking of target and masker speech streams, rD = rT-rM. (**D**) Average neural speech tracking across SI. (**E**) Averaged neural speech tracking across SNR. (**F**) rD across SNR for different groups of SI, with a linear line fitted to the data sample distribution of rD. Scatters and the fitted line are color-coded by SI. (**G**) Slope of rD vs. SNR with SI fixed. Significant slopes with 95% confidence intervals not containing 0 are marked with asterisks.

### 3.2. Increased SNR Leads to Reduced Attentional Effort and Neural Speech Tracking

Our findings of a negative relationship between SNR and neural speech tracking (rD) under high SI conditions suggest a secondary effect of SNR. One possibility is that increasing SNR may lead to decreased attentional performance (AP) and/or attentional effort (AE), which could consequently reduce rD. Specifically, AP is the is the actual performance outcome while AE refers to the cognitive resources required to maintain attention, including motivation and resource allocation (Pashler et al., 2001; Sarter et al., 2006). To investigate this, we used the hit rate of word repetition detection task (HR) as an ongoing measure of attentional performance, where a high HR indicates heightened attentional performance (Kirchner, 1958; Laffere et al., 2020; Marinato and Baldauf, 2019). Additionally, we used gaze velocity (GV), to measure oculomotor activity, as an indicator of ongoing attentional effort (AE); notably, a low GV suggests increased attentional effort (Ala et al., 2020; Ciccarelli et al., 2019; Gopher, 1973). (see **“4. Materials and Methods” - “2.4.3 Attention Measures”**). Fig 3A and 3B illustrate how SNR influences these two attention-related metrics. In Fig 3A we observe that the median HR levels off after -5 dB, suggesting that our attentional performance metric reaches a maximum beyond this SNR threshold, limiting its ability to explain changes in rD in this range.

**Fig 3.**
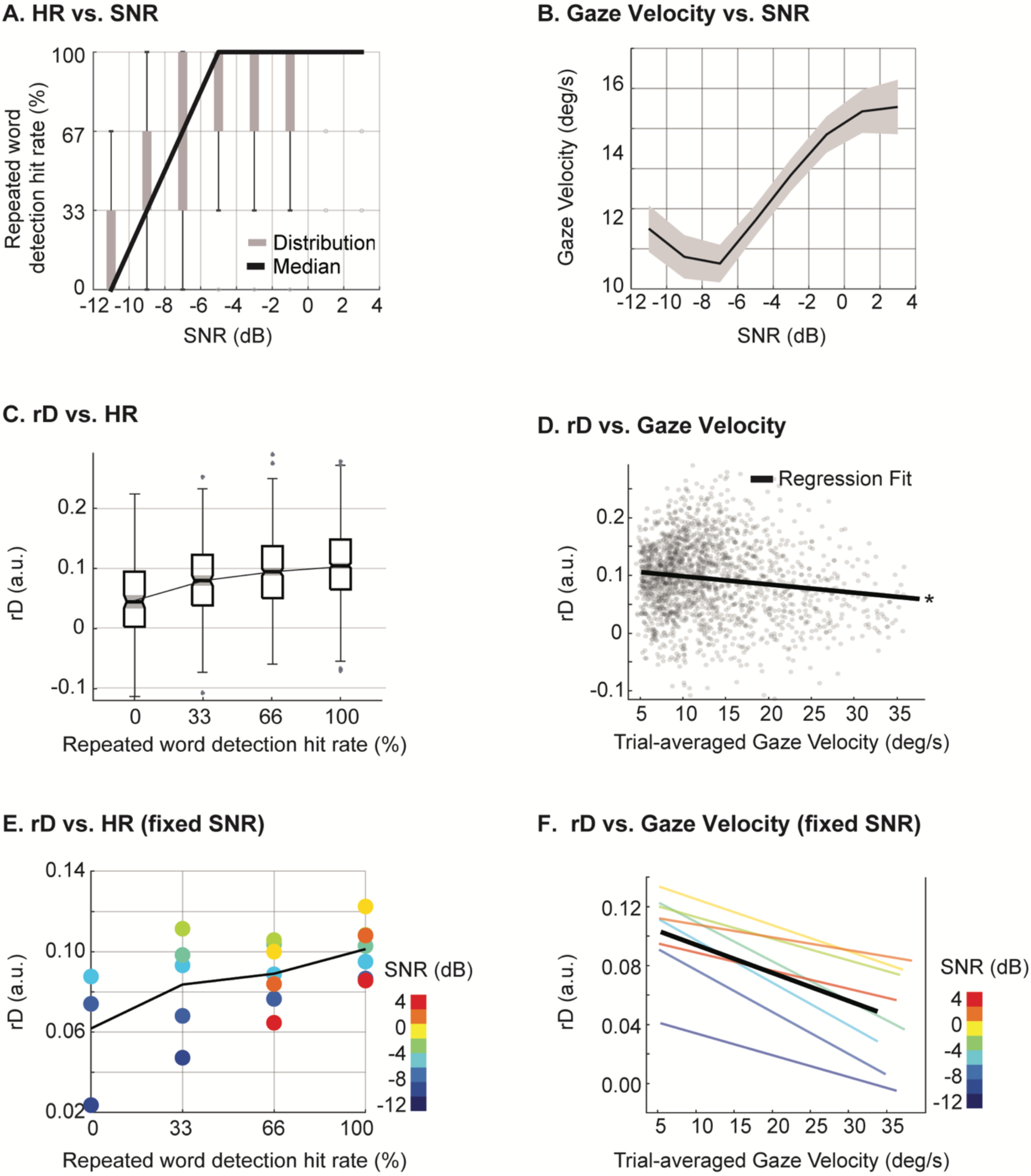
Attentional performance and attentional effort. (**A**) single-trial repeated word hit rate (HR) increases with SNR. (**B**) Gaze velocity (GV) increases with SNRs. (**C**) Interaction between rD and HR: distribution of rD across groups of HR. Medians and confidence intervals are marked with black lines and gray shades, respectively. Significant differences between groups are found (p<0.05, Kruskal-Wallis test, Bonferroni-corrected). (**D**) Interaction between rD and GV: GV negatively correlates with rD (c = -0.15, p<1e-5). The black line marks the regressed linear fit with significant slope and intercept: rD = -0.0014 * GV + 0.1126. (**E**) Interaction between rD and HR when fixing SNR. (**F**) Interaction between rD and GV when fixing SNR.

Conversely, Fig 3B shows that GV significantly increases at higher SNRs, where speech is highly intelligible. This indicates that the attentional effort required to maintain focus on the target talker, which is inversely related to GV, substantially decreases when SNR is sufficiently high. The reduced attentional effort, reflected by increased ocular activity, correlates with the decline in rD (Fig 3D, Pearson correlation test, c = -0.15, p<1e-5).

We conducted a more detailed analysis to explore the impact of attention-related features on rD. Fig 3C shows the modulation of rD by attentional performance, measured by HR. Higher HR was found to be associated with higher rD. Moreover, this change is not continuous due to the discrete nature of the behavioral response (three repeated words in each trial). The variation of rD with attentional effort, measured by GV, is shown in Fig 3D. This analysis reveals that trials with lower rD also exhibit more frequent gaze activity, suggesting that a decrease in attentional effort is correlated with decreased neural speech tracking.

To ensure these results apply across the entire range of SNR values, we repeated the same analyses separately for each SNR bin from -12 dB to 4 dB, as shown in Fig 3E and 3F. The results confirm that the observed positive correlation between rD and HR (Fig 3E) and the negative correlation between rD and GV (Fig 3F) is consistent across different SNR values. Hence, while the effect of attentional effort on rD becomes more visible in higher SNRs, these two show the same relationship in all SNR conditions. In summary, Fig 3 shows that attentional effort decreases with SNR, meaning subjects exert less effort in easier trials, which also corresponds to decreased target speech neural tracking.

### 3.3. Modeling the Interactions Between SNR, Speech Intelligibility, and Attentional Effort on Neural Speech Tracking

Given that our previous results indicate that multiple interacting variables influence rD, we used a computational model to elucidate these complex relationships. Specifically, we fitted a linear model to predict rD for each trial from that trial’s objective (SNR) and subjective (SI and GV) measurements (Adjusted *R*^/^: 0.151; F-statistic vs. constant model: 48.2, p < 0.001). The main effects and interaction terms are depicted in Fig 4A. This analysis shows that SI positively influences rD, while GV negatively affects rD. Interestingly, the direct influence of SNR on rD is not significant when interaction terms are included. This suggests an indirect influence of rD by SNR through the modulation of attentional effort and SI (Fig 4A). Specifically, increasing SNR improves SI and reduces attentional effort. The opposing effects of SI and attentional effort on rD could explain the non-linear relationship observed in Fig 2E.

**Fig 4.**
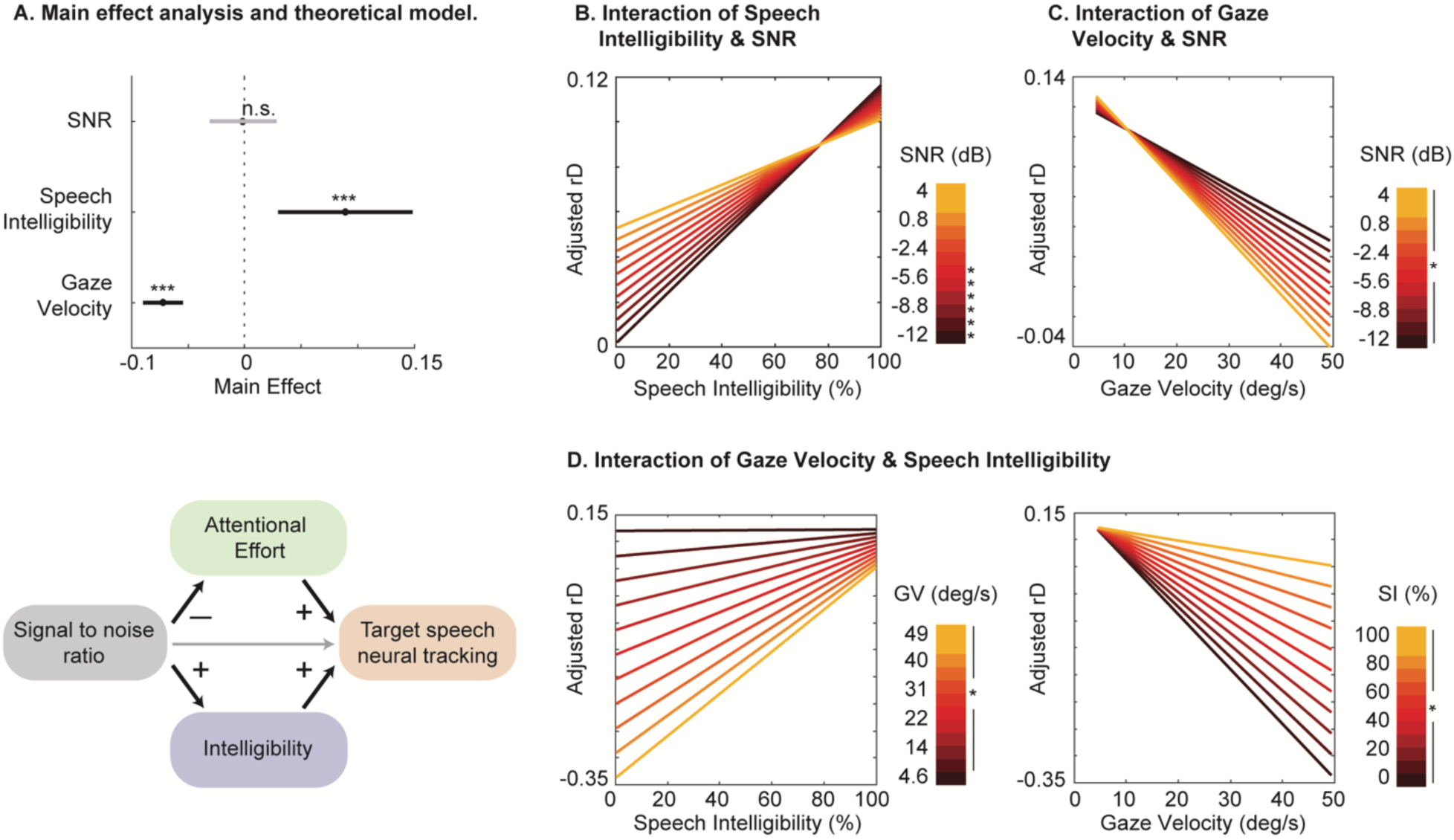
Interaction of various factors. (**A**). The main effect analysis of a linear model, and a hypothesized model of feature interactions. Speech intelligibility and gaze velocity exhibit a significant effect on rD (p<0.001, t-test), while SNR does not. (**B**) Interaction effect between SNR and SI of the fitted linear model. (**C**) Interaction effect between SNR and GV of the fitted linear model. (**D**) Interaction effect between SI and GV of the fitted linear model.

Fig 4B-4D further illustrate the interaction between these features. Fig 4B shows that SI has a positive impact on rD irrespective of SNR levels. However, as SNR increases, the effect of SI and its significance lessens, likely due to SI approaching the ceiling at higher SNRs. Fig 4C demonstrates the interaction between SNR and GV, showing that GV negatively impacts rD across all SNR levels. The change in the slope with increasing SNR suggests that rD is more sensitive to GV at higher SNRs. This increased sensitivity indicates that attentional effort plays a more significant role in shaping the neural target tracking at higher SNRs especially after maximum intelligibility.

Fig 4D shows how SI and GV differentially influence rD. Fig 4D (left) shows that the impact of SI on rD varies with different levels of attentional effort. With increased attentional effort (low GV), the influence of SI on rD decreases, highlighting the primary role of attention in shaping neural speech tracking. Conversely, Fig 4D (right) shows that the negative impact of increased GV on rD depends on SI. Attentional effort has the highest influence on rD in less intelligible conditions, where increased performance may attempt to compensate for the heightened difficulty of the listening task.

### 3.4 Temporal and Spatial Dynamics of Neural Responses Under Varying Speech Intelligibility and Attentional Effort

To investigate how the timing and spatial distribution of neural response patterns change under different SI and GV conditions, we calculated the temporal response functions (TRFs) (Ding and Simon, 2012b; Lalor et al., 2009) for target speech in different listening conditions. TRFs capture the brain’s temporal dynamics in response to continuous auditory stimuli, reflecting the relationship between the EEG signal and the speech envelope over lags at different electrodes. Normalized TRFs for target speech, averaged across all channels, are shown in Fig 5A and 5B.

**Fig 5.**
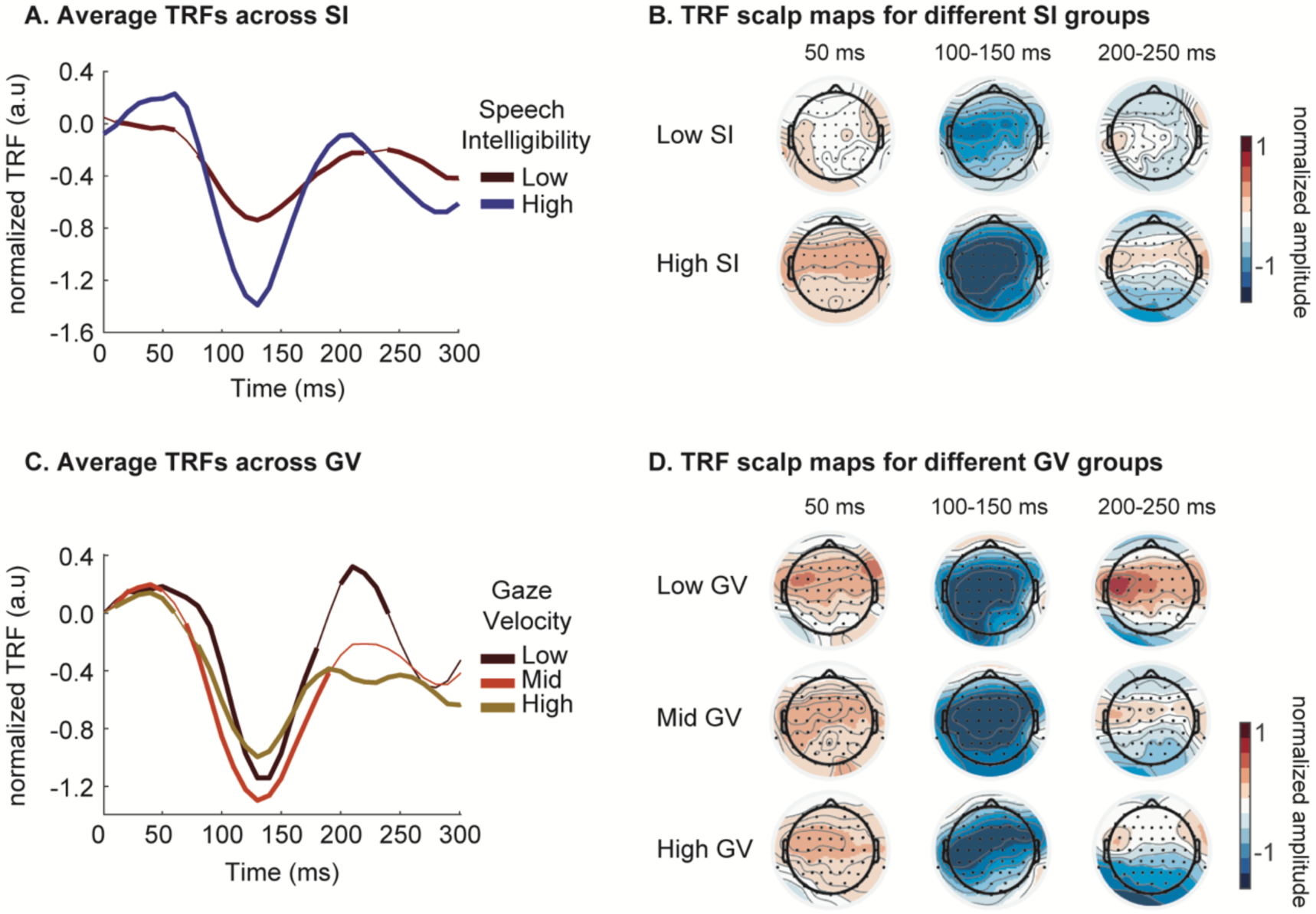
Normalized Temporal Response Functions (TRFs) under different Speech Intelligibility (SI) or Gaze Velocity (GV) levels. (**A**) The normalized TRFs under low (<50%) and high (>50%) SI. TRFs are averaged across all EEG channels. Significant temporal components (compared to the chance level, t-test, p<0.05) are marked with thick lines. (**B**) The topography of normalized TRFs amplitude at three critical time points (50ms, 100∼150ms, and 200∼250ms), under different levels of SI. (**C**) The normalized TRFs under low (<33%), mid (33%-67%), and high (>67%) GV. TRFs are averaged across all EEG channels. Significant temporal components (compared to the chance level, t-test, p<0.05) are marked in thick lines. (**D**) The topography of normalized TRFs amplitude at three critical time points (50ms, 100∼150ms, and 200∼250ms), under different levels of GV.

The group with low SI (< 50%) exhibits weaker early components TRF_50_ (positive, around 50 ms) across the scalp, especially in the temporal and central regions. Additionally, the low SI group shows reduced attention-related TRF components TRF_100_ (negative, around 100 ms) and TRF_200_ (positive, around 200 ms), indicating reduced selectivity for target speech and less suppression of masker speech components (Ding and Simon, 2012b; Fiedler et al., 2019).

Attentional effort, measured inversely by GV, also impacts the TRFs. While the acoustics-modulated early components TRF_50_ remain consistent across different GV, a significant difference emerges for higher-level, attention-related components around TRF_100_ and TRF_200_ (Fig 5C and 5D). In trials with lower attentional effort (high GV, GV>66.7% percentile), TRF_100_ responses decrease in posterior electrodes (Fig 5D). For TRF_200_, only the group with low GV (GV<33.3 % percentile) shows strong activation in the anterior and central areas.

## 4. Discussion

We demonstrate that neural tracking of target speech is influenced by both objective (signal to noise ratio) and subjective (speech intelligibility, attentional performance and effort) factors in distinct ways. As speech intelligibility increases, the positive effect of improving SNR on neural tracking of target speech diminishes. Specifically, in conditions where speech is highly intelligible, further increases in SNR decrease neural speech tracking. We propose that this decrease is caused by the reduced attentional effort required to focus on the target speech. Our findings show that gaze velocity, a measure proposed for quantifying attentional effort, effectively explains this reduction in neural speech tracking accuracy. Together, our findings suggest a complex interaction between speech intelligibility and attentional effort mediated by SNR in shaping the neural representation of speech in noise.

### 4.1. Distinct Contributions of SI and SNR to Neural Speech Tracking

Despite the significant correlation between speech intelligibility (SI) and signal-to-noise ratio (SNR), our findings reveal distinct impacts of each on neural tracking of target speech as measured by EEG. While SI and SNR are often discussed together due to their high correlation, they affect neural speech tracking through different mechanisms. Previous studies, including those by (Das et al., 2018; Iotzov and Parra, 2019; Vanthornhout et al., 2018), primarily investigated the impact of SI on neural tracking at different SNRs. Our results, however, underscore the importance of differentiating the effects of SI and SNR.

Objective acoustic features like SNR drive bottom-up processing in speech perception. In contrast, SI is shaped by an individual’s bottom-up perception and top-down processing capabilities and strategies for allocating cognitive resources or restoring features masked by noise (Raghavan et al., 2023). This distinction highlights how SNR and SI contribute differently to neural processing. For example, the activity in brain regions responsible for top-down processing is increased when bottom-up processing was impaired by degraded speech (lower SNRs) (Zekveld et al., 2006). Several previous studies also suggested different modulation of neural speech tracking by objective or perceptual speech attributes. Previous research has shown that while SNR non-linearly modulates the amplitude of the temporal response function, changes in neural latency align more closely with variations in SI (Yasmin et al., 2023). In contrast, the relationship between neural speech tracking and SI does not exhibit such non-linearity (Decruy et al., 2020a). Instead, the different metrics of neural speech tracking accuracy show slightly dissimilar correlations with SNR and SI, but this mismatch has not been explained (Nogueira and Dolhopiatenko, 2022). Our findings go beyond these observations by dissociating the interplay between SI and SNR and their impact on neural speech tracking. We provide evidence that SNR contributes indirectly to neural speech tracking by modulating attentional effort and speech intelligibility. This supports the notion of an indirect contribution of SNR to neural speech tracking, as suggested by (Etard and Reichenbach, 2019). It is important to note that our measure of neural speech tracking is based on EEG recordings, which reflect scalp potentials from large populations of neurons but do not provide the fine-grained detail available from invasive or single-neuron recordings. Studies using invasive techniques in animals and humans have shown that noise-invariant representations gradually develop along the auditory pathway (Kell and McDermott, 2019; Mesgarani et al., 2014; Rabinowitz et al., 2013), with lower areas representing the noise and higher areas filtering it out. Our findings highlight how the combined effects of these interactions manifest in scalp EEG signals which is critical as EEG is the most widely used measure to study speech in noise in normal hearing, hearing impaired, and aging individuals (Di Liberto et al., 2022; Fuglsang et al., 2020; Mesik et al., 2021).

### 4.2. The Role of Attention in Neural Speech Tracking

Attention plays a crucial role in how the brain tracks attended speech. The acoustic characteristics of speech can influence attention levels and, consequently, the accuracy of neural speech tracking (Ding and Simon, 2012a; Iotzov and Parra, 2019; Mesgarani and Chang, 2012; Power et al., 2012; Vanthornhout et al., 2019a; Zion Golumbic et al., 2013). Our study examined two aspects of attention: attentional performance and attentional effort. Attentional effort refers to the cognitive resources required to maintain attention, including motivation and resource allocation, while attentional performance is the actual outcome (Pashler et al., 2001; Sarter et al., 2006). We assessed attentional performance using repeated word hit rate (HR). We inferred attentional effort by measuring gaze velocity (GV) (Ala et al., 2020; Ciccarelli et al., 2019; Gopher, 1973). In easier listening conditions (SNR > -1.6 dB), we observed a significant reduction in neural speech tracking accuracy with increasing SNR. This finding aligns with studies by (Das et al., 2018; Lesenfants et al., 2019), which noted decreased neural speech tracking accuracy from mildly noisy to clean conditions. Our study further investigates this paradoxical relationship by measuring ocular activity as an approximation of attentional effort. The identified negative interaction between ocular activity and task difficulty was also illustrated by (Contadini-Wright et al., 2023; Cui and Herrmann, 2023; Herrmann and Ryan, 2024). In contrast, attentional performance, measured by HR, shows a limited correlation with task difficulty. These results suggest that the reduction in neural target speech tracking can be more accurately attributed to changes in attentional effort rather than variations in attentional performance. Our findings also align with previous research indicating that increased eye movement activity reflects less suppression of task-irrelevant psychological activity, impairing information processing such as selective neural speech perception (Abeles et al., 2020; Braga et al., 2016; Cui and Herrmann, 2023). More importantly, we provide a potential explanation for the reduced neural speech tracking in easier listening conditions, as also reported by (Das et al., 2018; Hauswald et al., 2022; Lesenfants et al., 2019). Note that this decreasing effect exists across SNRs, not only in the high SNR listening conditions. Attentional effort and attentional performance exhibit distinct characteristics despite their interconnectedness (Bruya and Tang, 2018). Our study also supports differentiating the modulation of neural entrainment between attentional effort and attentional performance, similar to the findings of (Dai and Shinn-Cunningham, 2016), which showed that selective attention could modulate the strength of cortical event-related potential but not change the attentional performance. It is also worth mentioning studies that have demonstrated increased neural speech tracking in older populations and subjects with hearing impairment (Decruy et al., 2020b, 2019). Our study offers an explanation for these observations: increased task difficulty in these subject populations elevates attentional effort, thereby enhancing neural speech tracking. To test this hypothesis, we propose measuring differences in gaze velocity between populations or adjusting the SNR to identify the threshold at which neural speech tracking declines relative to normal hearing subjects, estimated in our study at approximately -1.6 dB.

### 4.3 Modeling the Interplay of Speech Intelligibility and Attentional Effort on Neural Speech Tracking

In analyzing the interaction of various features on predicting neural speech tracking, we found that both speech intelligibility (SI) and gaze velocity (GV) have significant effects, while signal-to-noise ratio (SNR) does not. Supplementary analyses and comparisons of temporal response functions (TRFs) for significant features (SI and GV) revealed that SI influences both acoustic-related (TRF_50_) (Ding and Simon, 2013) and attention-related components (TRF_100_, TRF_200_) (Ding and Simon, 2012b; Fiedler et al., 2019) of neural speech tracking, consistent with previous studies (Chen et al., 2023; Muncke et al., 2022). Notably, the modulation of early response (TRF_50_) may be attributed to the combined effect of SNR and SI, as shown in prior study, where a lower SNR at the same SI resulted in reduced TRF_50_ amplitude (Verschueren et al., 2020). In contrast, GV, as an indicator of attentional effort, only modulates attention-related components, specifically the activation area of TRF_100_ and the intensity of TRF_200_. These components are closely associated with the top-down process of directing mental resources toward the target of interest (Fritz et al., 2007; Kong et al., 2014; Vanthornhout et al., 2019b). These findings suggest that while SI affects multiple aspects of neural speech processing, GV’s influence is limited to the attentional mechanisms.

From the detailed analyses of the interaction among features, we proposed a model based on our finding that AE and SI show counterbalancing effect on neural speech tracking as SNR increases, with the dominant factor shifting from SI to AE. The proposed model is able to explain the widely observed non-linearity between task demands and neural speech tracking (Das et al., 2018; Hauswald et al., 2022; Lesenfants et al., 2019), and also provides an explanation for the increased speech tracking in hard of hearing and aging populations (Decruy et al., 2020b, 2019), for which the increased task difficulty results in an increased attentional effort.

There are several limitations to consider while interpreting our results. As EEG signals provide only a broad overview of cortical activity, complementary neuroimaging techniques would be needed to fully characterize the encoding of noisy speech in various cortical and subcortical auditory regions. Additionally, our measure of attentional effort is indirect. While used extensively in the field (Ala et al., 2020; Ciccarelli et al., 2019; Gopher, 1973), gaze velocity is only an approximation of the cognitive resources that are used to maintain focus. Finally, our measure of attentional performance is sparse, as we cannot rule out the possibility that the listeners lose focus in between the repeated words. Future research is needed to explore more direct methods to measure cognitive load and attentional performance, and to expand these findings to aging and hard of hearing population.

In summary, our study demonstrates that the neural tracking of target speech is influenced by SNR, speech intelligibility, and attentional performance and attentional effort, with distinct and sometimes opposing effects. By disentangling the roles of attentional performance and effort, we provide a clearer understanding of how these factors interact to shape neural speech processing. Beyond their scientific impact, these insights also have important implications for developing auditory technologies and strategies to improve speech perception in noisy environments.

## Author Contributions

XH, VR, and NM conceived the project. XH and NM designed the experiment and analyzed the data. XH and NM wrote the manuscript, and all authors provided feedback and revisions.

